# Hyperactive mTORC1/4EBP1 Signaling Dysregulates Proteostasis and Accelerates Cardiac Aging

**DOI:** 10.1101/2024.05.13.594044

**Authors:** Weronika Zarzycka, Kamil A Kobak, Catherine J King, Frederick F Peelor, Benjamin F Miller, Ying Ann Chiao

**Author notes:** **Corresponding Author:** Ying Ann Chiao.

## Abstract

The mechanistic target of rapamycin complex 1 (mTORC1) has a major impact on aging by regulation of proteostasis. It is well established that mTORC1 signaling is hyperactivated with aging and age-related diseases. Previous studies have shown that partial inhibition of mTOR signaling by rapamycin reverses the age-related decline in cardiac function and structure in old mice. However, the downstream signaling pathways involved in this protection against cardiac aging have not been established. TORC1 phosphorylates 4E-binding protein 1 (4EBP1) to promote the initiation of cap-dependent translation. The aim of this project is to examine the role of the mTORC1/4EBP1 axis in age-related cardiac dysfunction. We utilized a whole-body 4EBP1 KO mouse model, which mimics a hyperactive 4EBP1/eIF4E axis, to investigate the effects of hyperactive mTORC1/4EBP1 axis in cardiac aging. Echocardiographic measurements revealed that young 4EBP1 KO mice have no difference in cardiac function at baseline compared to WT mice. Interestingly, middle-aged (14–15-month-old) 4EBP1 KO mice show impaired diastolic function and myocardial performance compared to age-matched WT mice and their diastolic function and myocardial performance are at similar levels as 24-month-old WT mice, suggesting that 4EBP1 KO mice experience accelerated cardiac aging. Old 4EBP1 KO mice show further declines in systolic and diastolic function compared to middle-aged 4EBP1 KO mice and have worse systolic and diastolic function than age-matched old WT mice. Gene expression levels of heart failure markers are not different between 4EBP1 KO and WT mice at these advanced ages. However, ribosomal biogenesis and overall protein ubiquitination are significantly increased in 4EBP1 KO mice when compared to WT, which suggests dysregulated proteostasis. Together, these results show that a hyperactive 4EBP1/eIF4E axis accelerates cardiac aging, potentially by dysregulating proteostasis.

## Introduction

Cardiovascular disease (CVD) is a predominant cause of death worldwide [1, 2]. According to the 2022 update of the American Heart Association report, 27% of all global deaths were caused by CVD [2]. Moreover, prevalence of CVDs arise with age because aging itself is a primary risk factor for CVDs [1]. Adults over 70 years of age constitute the majority of the population suffering from CVD [1]. Aging is associated with a progressive healthspan decline and deterioration of the physiological functions at whole organism and individual organ levels, which in turn leads to age-related diseases and ultimately to death [3]. Therefore, it is not surprising that cardiac aging causes deterioration of heart function, and increased prevalence and susceptibility to heart failure [1]. Cardiac aging is defined as an age-dependent progressive degeneration, which makes the heart and vessels more vulnerable to stress, contributing to increased mortality and morbidity [4-6]. The Framingham Heart Study (FHS) and The Baltimore Longitudinal Study on Aging (BLSA) established the key features of cardiac aging phenotype [5, 6]. Of these, diastolic dysfunction is one of the most significant impairments in the aged heart, as well as prolonged isovolumic relaxation and isovolumic contraction. Conversely, systolic function is relatively preserved and only slightly declines with age [7, 8]. In terms of structural changes, aging also results in left ventricular, left atrial hypertrophy and increased arterial stiffness [5-7]. Due to the progressive aging of the population, the burden of age-related CVD and cardiac dysfunction in the US and worldwide is increasing. Therefore, it is important to study the molecular mechanisms underlying the age-related deterioration of cardiac structure and function to prevent the increasing prevalence of CVD.

López-Otin et al. described nine hallmarks of aging by categorizing molecular and cellular changes that happen with age [9, 10]. Dysregulated nutrient sensing and loss of proteostasis are two hallmarks of aging [9]. The mechanistic target of rapamycin complex 1 (mTORC1) is nutrient-sensing system that is responsible for the regulation of many essential anabolic aspects of metabolism [9, 11]. It responds to and is activated by high amino acid concentrations and excess nutrients to promote protein synthesis and inhibit autophagy [9, 11, 12]. mTOR plays a pivotal role in aging and longevity [11]. It has been shown that inhibition of mTOR signaling by caloric restriction (CR) or mTOR inhibitor rapamycin results in lifespan extension in model organisms such as yeast, worms, flies and mice [13-17]. Moreover, we and others have also shown that partial inhibition of mTOR signaling by rapamycin reverses the age-related deteriorations in cardiac function and structure in old mice [18-20].

mTOR signaling, due to its role in protein synthesis and degradation, is crucial for maintenance of protein homeostasis (proteostasis). Impaired proteostasis has been demonstrated as underlying mechanisms of aging and age-related diseases [21-24]. It is manifested by impaired translation, accumulation of misfolded proteins, protein aggregation or enhanced ubiquitination [21-24]. Impaired function of protein quality control mechanism such as unfolded protein response (UPR) can also lead to impaired proteostasis maintenance [25]. Based on previous studies, protein quality and age-related proteostasis imbalance are established factors contributing to cardiac diseases [26-29]. mTORC1 regulates proteostasis through its three key downstream effectors: Unc-51 Like Autophagy Activating Kinase 1 (ULK1), S6 kinase 1 (S6K1), and 4E-binding protein 1 (4EBP1) [30-32]. ULK1 governs protein degradation via autophagy [32]. When mTORC1 is active, it phosphorylates ULK1 resulting in autophagy inhibition [32]. S6K1 and 4EBP1, on the other hand, control protein synthesis [31, 33]. mTORC1 phosphorylates 4EBP1, which suppresses its binding to eukaryotic initiation factor 4E (eIF4E), releasing eIF4E for the initiation of cap-dependent translation [30]. Sonenberg’s group has shown that cardiac specific knockout of mTOR is lethal and leads to cardiac hypertrophy and heart failure [34]. Moreover, they showed that cardiac specific mTOR knockout (Mtor-cKO) mice have higher expression and increased dephosphorylation of 4EBP1 along with cardiomyocytes apoptosis compared to control mice [34]. Interestingly, whole-body deletion of 4EBP1 improved survival of Mtor-cKO mice and reduced cardiomyocytes apoptosis, strongly suggesting an important role of mTOR/4EBP1 axis for cardiac cell survival and function [34]. Contrarily, deletion of 4EBP1 in *D. melanogaster* led to lifespan shortening and abolition of lifespan extension induced by dietary restriction [11, 35]. Moreover, Robert Wessells et al. showed that in *Drosophila*, overexpression of 4EBP prevented the age-related decline in cardiac function while deletion of 4EBP lead to a high failure rate at young age comparing to revertant controls [36]. Despite these significant strides in understanding the roles of mTOR signaling and its downstream effectors, the role of 4EBP1 in regulating cardiac aging in mammalian models remains elusive. Inhibition of mTOR signaling by rapamycin or CR reverses cardiac aging in mice [18], however, the involvement of 4EBP1 and other mTOR downstream targets in cardiac aging have not been established.

The current study investigates the role of the mTORC1/4EBP1 signaling axis in age-related cardiac dysfunction. We hypothesized that hyperactive mTORC1/4EBP1 signaling impairs proteostasis maintenance and augments age-related cardiac dysfunction in mice. We employed a whole body 4EBP1 KO mouse model to study the effects of a hyperactive 4EBP1 axis in the hearts at different ages. We found that 4EBP1 KO mice show accelerated cardiac aging phenotype comparing to WT controls. Moreover, 4EBP1 KO mouse hearts have dysregulated proteostasis. A better understanding of the molecular mechanisms underlying cardiac aging is crucial for developing interventions for early prevention and effective treatment of cardiovascular disease in the aging population.

## Methods

### Animals

Animals handling and all procedures were approved and performed according to the Institutional Animal Care and Use Committee (IACUC) at Oklahoma Medical Research Foundation. Whole-body homozygous 4EBP1 knockout (KO) mouse line, generated and derived from C57BL/6 strain, was kindly provided by Dr. Nahum Sonenberg at McGill University [37]. To study age-dependent cardiac dysfunction, male and female mice of three age groups were used: young (3–7-month-old), middle-aged (14–16-month-old) and old (24-month-old).

### Echocardiography

Cardiac function was assessed by echocardiography using ACUSON CV70 Ultrasound Machine (Siemens) with a 13-5 MHz transducer as previously described [18]. Prior to measurements mice were anesthetized with isoflurane and their heart rate was maintained at approximately 400-500 beats per minute. Systolic function was assessed by M-mode images. Fractional shortening together with left ventricular internal diameter at diastole (LVIDd) and left ventricular posterior wall thickness at diastole (LVPWd) were measured. Parameter of diastolic function (Ea/Aa) was measured by Tissue Doppler imaging at the mitral annulus. From pulse-wave Doppler imaging the sum of isovolumetric contraction time (IVCT) and isovolumetric relaxation time (IVRT) was divided by ejection time (ET) to calculate the myocardial performance index (MPI).

### Tissue Harvest and Processing

Prior to sacrifice, mice were subjected to 4-hour fasting. Mice were anesthetized using isoflurane and weighed. Rib cage was open to expose the heart and blood was collected by cardiac puncture. Heart tissue was weighed and immediately frozen with liquid nitrogen. Tibia length was measured to calculate heart weight (HW) to tibia length (TL) ratio to determine normalized heart weight and cardiac hypertrophy. Wet and dry lung weights were measured to assess pulmonary edema. Frozen cardiac tissue was pulverized using Qiagen TissueLyser II for downstream molecular analyses.

### Analysis of Gene Expression Levels

Total RNA was isolated from frozen heart tissue powder using Qiagen RNeasy Fibrous Tissue Mini Kit (Cat. No. / ID: 74704). Using nanodrop RNA concentration was measured. Reverse transcription reaction was conducted to obtain cDNA using 1 μg RNA and Maxima FirstStrand cDNA Synthesis kit (Thermo Fisher Scientific). Gene expression levels were quantified by performing quantitative real-time PCR using TaqMan Gene Expression Assays (Thermo Fisher Scientific) with CFX96 Touch Real-Time PCR Detection System (Bio-Rad). Life Technologies TaqMan probes: Atf4 (Mm00515325_g1), Gdf15 (Mm00442228_m1) Mb (Mm00442968_m1), Myh6 (Mm00440359_m1), Myh7 (Mm00600555_m1), Nppa (Mm01255747_g1), Nppb (Mm01255770_g1) were used for gene expression analyses.

### Western Blotting Assay

Proteins were extracted from pulverized hearts using 1x RIPA Buffer with addition of protease and phosphatase inhibitor cocktail (HALT, Thermo Fisher Scientific) and deacetylase inhibitors (10 μM Trichostatin A (Sigma) and 10mM nicotinamide (Thermo Fisher Scientific)). Protein concentrations were measured by using bicinchoninic acid assay (BCA, Thermo Fisher Scientific). Protein samples were loaded on SDS-PAGE Criterion precast 4-15% gradient gel using Criterion System (Bio-Rad). Resolved proteins were transferred onto PVDF membrane by using Criterion Blotter system (Bio-Rad). Total protein staining was performed by using Revert 700 Total Protein Stain (LICOR). After removal of Revert stain, the membrane was blocked with 5% BSA in TBST and incubated with primary antibody in 5% BSA in TBST. Following primary antibodies were used for Western blot analysis: 4EBP1(1:1000, #9452 – Cell Signaling), Troponin I (1:1000, #4002 – Cell Signaling), pSer23/24 Troponin I (1:1000, #4004S – Cell Signaling), pSer150 Troponin I (1:1000, #PA535410 - Invitrogen), Ubiquitin (1:1000, #58395S – Cell Signaling). Membrane was incubated with secondary antibody conjugated with HRP and proteins were visualized by using chemiluminescence assay (Pierce) and G:BOX Syngene Imaging System. Proteins were quantified by densitometry using AlphaView (ProteinSimple).

### Deuterium Oxide Labelling and Isotope Analysis

Middle-aged WT and 4EBP1 KO mice (n=5) were intraperitoneally injected with the bolus dose of 99% deuterium oxide (D_2_O) with 0.9% sodium chloride to initiate labelling at 30 μl/g body weight. For subsequent 15 days 8% D_2_O enriched drinking water was given to mice. After 15-day labelling period mice were euthanized and tissues were collected as above-mentioned. Protein and RNA synthesis were analyzed according to our previously published procedures[38, 39]. For measurement of protein synthesis approximately 30 mg of pulverized whole heart tissue was homogenized in isolation buffer (100 mM KCl, 40 mM Tris–HCl, 10 mM Tris Base, 5 mM MgCl2, 1 mM EDTA, 1 mM ATP, pH = 7.6) with addition of phosphatase and protease inhibitors (HALT, Thermo Fisher Scientific). Next, by differential centrifugation subcellular fractions were isolated to obtain myofibrillar, mitochondrial and collagen protein fractions. Individual protein fractions were next derivatized for subsequent analysis of deuterium enrichment in alanine using Gas Chromatography-Mass Spectroscopy (Agilent 7890B GC coupled to Agilent 5977B MS, Agilent, Santa Clara, CA). To assess ribosomal biogenesis total RNA was extracted from ∼10 mg of heart tissue using Qiagen RNeasy Fibrous Tissue Mini Kit (Qiagen). Concentration and quality of isolated RNA were determined by NanoDrop (Thermo Fisher Scientific). Next, RNA was hydrolyzed, derivatized, and analyzed using GC-MS (Agilent 8890GC coupled to Agilent 7010B MS, Agilent, Santa Clara, CA). MassHunter software was used to analyze MS data [39].

To determine body water enrichment, serum plasma was distillated overnight at 80°C, diluted 1:300 in dd H2O and analyzed on a liquid water isotope analyzer (Los Gatos Research, Los Gatos, CA, USA) against a standard curve prepared with samples containing different concentrations of D_2_O.

To obtain protein and RNA synthesis rates, we used standard FSR calculations as previously published [39, 40]. Briefly, the newly synthesized fraction (f) of proteins was calculated from the enrichment of protein-bound alanine, divided by the true precursor enrichment (p), using plasma D_2_O enrichment with Mass Isotopomer Distribution Analysis (MIDA) [41]. RNA synthesis was determined by deuterium incorporation into purine ribose of RNA as previously published [39, 40] with MIDA adjustment of the equilibration of the enrichment of the body water pool with purine ribose.

### Statistical Analysis

GraphPad Prism 9.0 was used to perform all the data analysis. Unpaired two-tailed Student’s t-test was used for experiments involving two groups. Ordinary one-way ANOVA was used for experiments involving more than two groups, using Šídák’s correction for multiple comparisons. All data are expressed as mean ± SEM, p< 0.05 was considered significant.

## Results

### 4EBP1 deficiency leads to cardiac dysfunction in middle-aged and old, but not young, mice

Previous study has shown that deletion of ribosomal S6 kinase (S6K), one of the mTORC1 downstream pathways, does not have an impact on development of neither pathological nor physiological cardiac hypertrophy [42]. However, the role of 4EBP1 downstream mTORC1 effector in regulation of cardiac function has not been established in mammals yet. To determine the involvement of mTORC1/4EBP1 axis in cardiac aging, we examined cardiac function and structure of young (3-7 months), middle-aged (14-16 months) and old (24 months) whole-body 4EBP1 knockout (KO, Fig. 1A) male mice by echocardiography. Young 4EBP1 KO mice show no cardiac hypertrophy nor pulmonary edema (Fig. 1B, C). Cardiac systolic (fractional shortening, FS) and diastolic function (Ea/Aa) and myocardial performance (myocardial performance index, MPI) of young 4EBP1 KO mice are not different when compared to young C57BL/6 wild-type (WT) mice (Fig. 1D-F). However, we found that middle-aged and old 4EBP1 KO mice have worse diastolic function than age-matched WT controls, and diastolic function of middle-aged 4EBP1 KO mice is similar to that of old WT mice (Fig. 2A). Middle-aged 4EBP1 KO mice show a trend of impaired myocardial performance (represented by an increase in MPI) comparing to old WT (Fig. 2B). Middle-aged and old 4EBP1 KO mice have significantly worse systolic function when compared to age-matched control mice (Fig. 2C), indicating that 4EBP1 deletion leads to systolic dysfunction in advanced ages. The impaired cardiac function of 4EBP1 KO mice in advanced ages, but not at young age, suggests an important role of mTORC1/4EBP1 axis in promoting age-related cardiac dysfunction.

**Figure 1.**
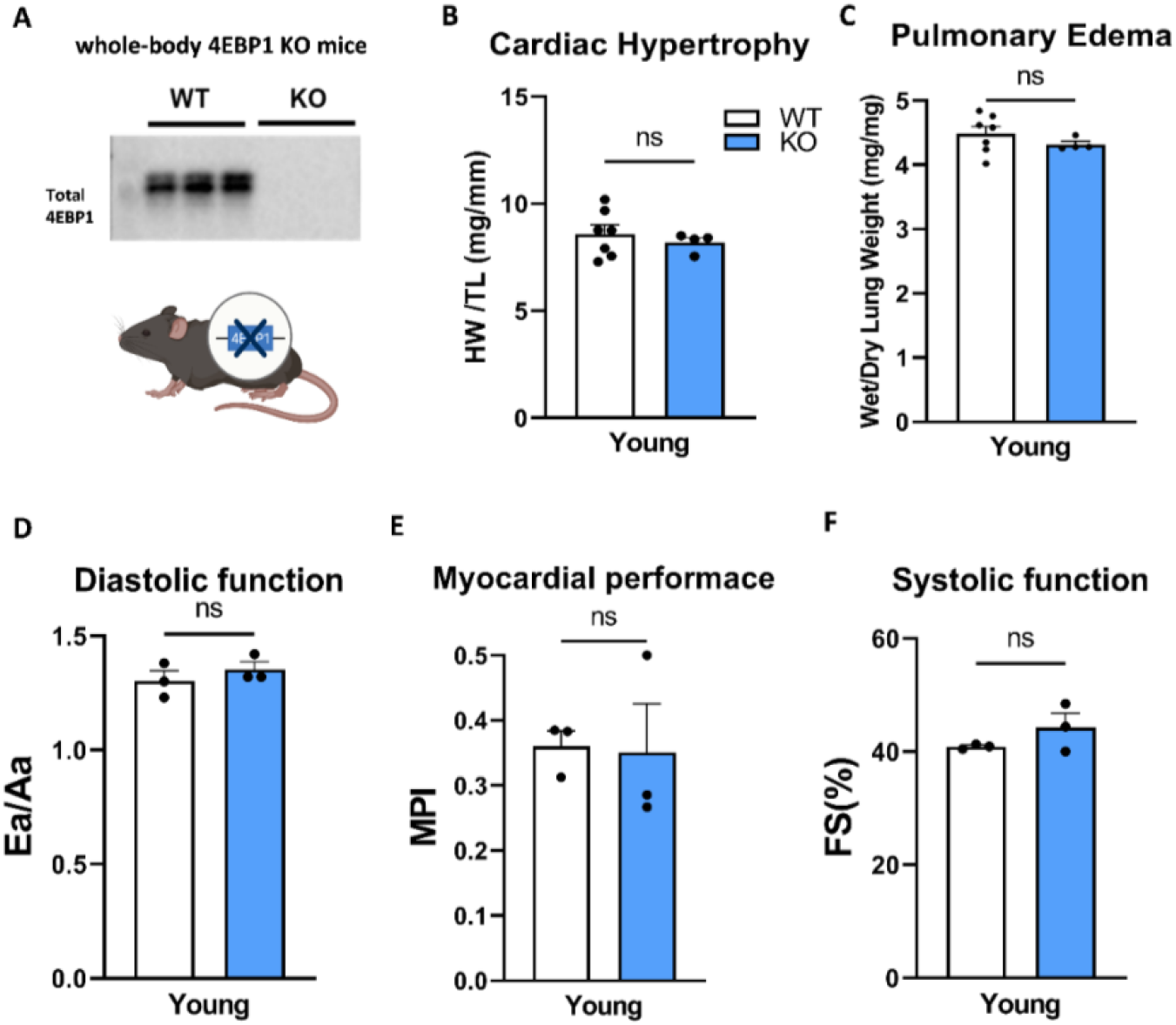
Complete 4EBP1 deficiency does not have an impact on cardiac hypertrophy and pulmonary edema and does not induce cardiac dysfunction in young male mice. **(A)** Western blot confirmation of whole-body 4EBP1 knockout mouse model. **(B)** Cardiac hypertrophy nor **(C)** pulmonary edema are different between young WT and 4EBP1 KO male mice. **(D)** Diastolic function (ratio of peak early to late diastolic mitral annular velocity, Ea/Aa), **(E)** myocardial performance (myocardial performance index, MPI) and **(F)** systolic function (fractional shortening, FS) are not different between young WT and 4EBP1 KO male mice, indicating no cardiac dysfunction. Values represent means ± SEM. Unpaired two-tailed Students t-test was used, n=3-7.

**Figure 2.**
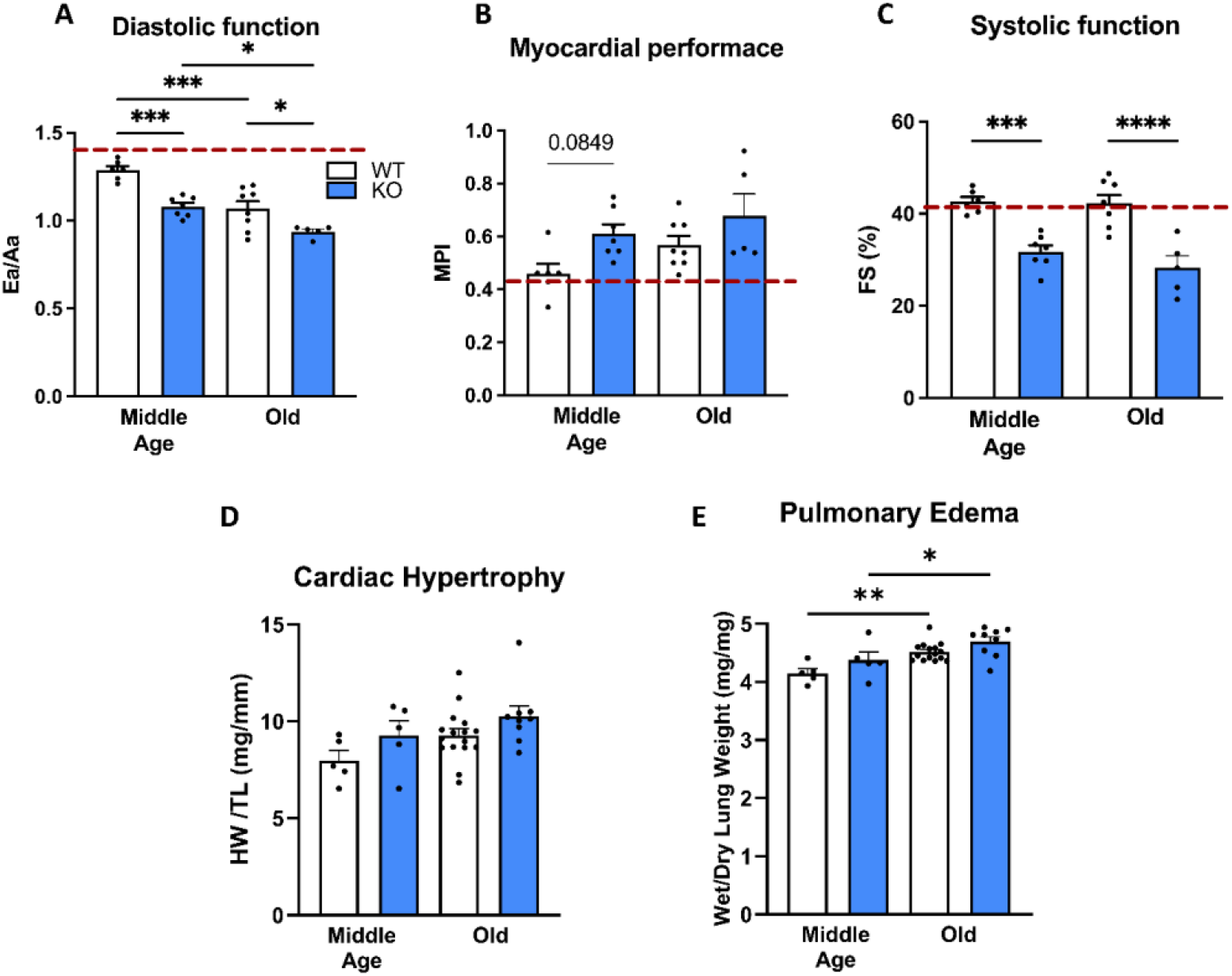
Deletion of 4EBP1 protein exacerbates cardiac dysfunction in mice without affecting cardiac hypertrophy and pulmonary edema. **(A)** Middle-aged (14-15-month-old) 4EBP1 KO mice have impaired diastolic function (Ea/Aa) and **(B)** myocardial performance index (MPI) at levels similar to 24-month-old WT mice. All these parameters further worsen in old KO mice. **(C)** Middle-aged and old 4EBP1 KO mice have worse systolic function than WT mice when fractional shortening (FS) was measured. Red dotted line indicates baseline measurement for young control mice. **(D, E)** Cardiac hypertrophy and pulmonary edema were not affected by 4EBP1 KO. Although, impact of age on cardiac hypertrophy resulting in increased hypertrophy with age was observed. Values represent means ± SEM. *p< 0.05; **p< 0.001; ***p< 0.0001 **(A-E)** Ordinary one-way analysis of variance (ANOVA) was performed, n=5-15.

Although, 4EBP1 KO male mice have worse cardiac function at advanced ages, HW/TL and wet/dry lung weight are not different from WT mice regardless of age and sex (Fig. 1B-C, Fig. 2D-E, Supplemental Fig. 1A, B). This indicates that mTORC1/4EBP1 axis does not affect cardiac hypertrophy nor pulmonary edema.

### 4EBP1 deletion does not alter gene expression levels of hypertrophy and senescence markers and phosphorylation of Troponin I

To investigate molecular changes caused by 4EBP1 deletion at advanced ages, we measured gene expression levels of hypertrophy and heart failure markers including natriuretic peptide b (Nppb), natriuretic peptide a (Nppa), myosin heavy chain 7 to 6 ratio (Myh7/Myh6) and growth differentiation factor 15 (Gdf15) (Fig. 3 A-D). We did not observe any significant changes in expression levels of above-mentioned genes upon 4EBP1 deletion. Activating transcription factor 4 (Atf4) is a regulator of the integrated stress response and has been shown as a downstream target of 4EBPs mediated translational regulation [12]. We measured expression levels of Atf4 and found no difference in Atf4 expression between WT and 4EBP1 KO mice (Supplemental Fig. 2A). Cellular senescence of cardiac cells has been shown as a contributing mechanism of age-related cardiac dysfunction [43, 44]. We examined expression of senescence markers, IL6, p16 and p19 and observed significantly increased expression of p16 with age (Supplement Fig. 2C). However, none of these genes show differences in transcript levels upon deletion of 4EBP1 (Supplement Fig. 2B-D). Similar results were also observed in female mice (Supplemental Figure 1C-F).

**Figure 3.**
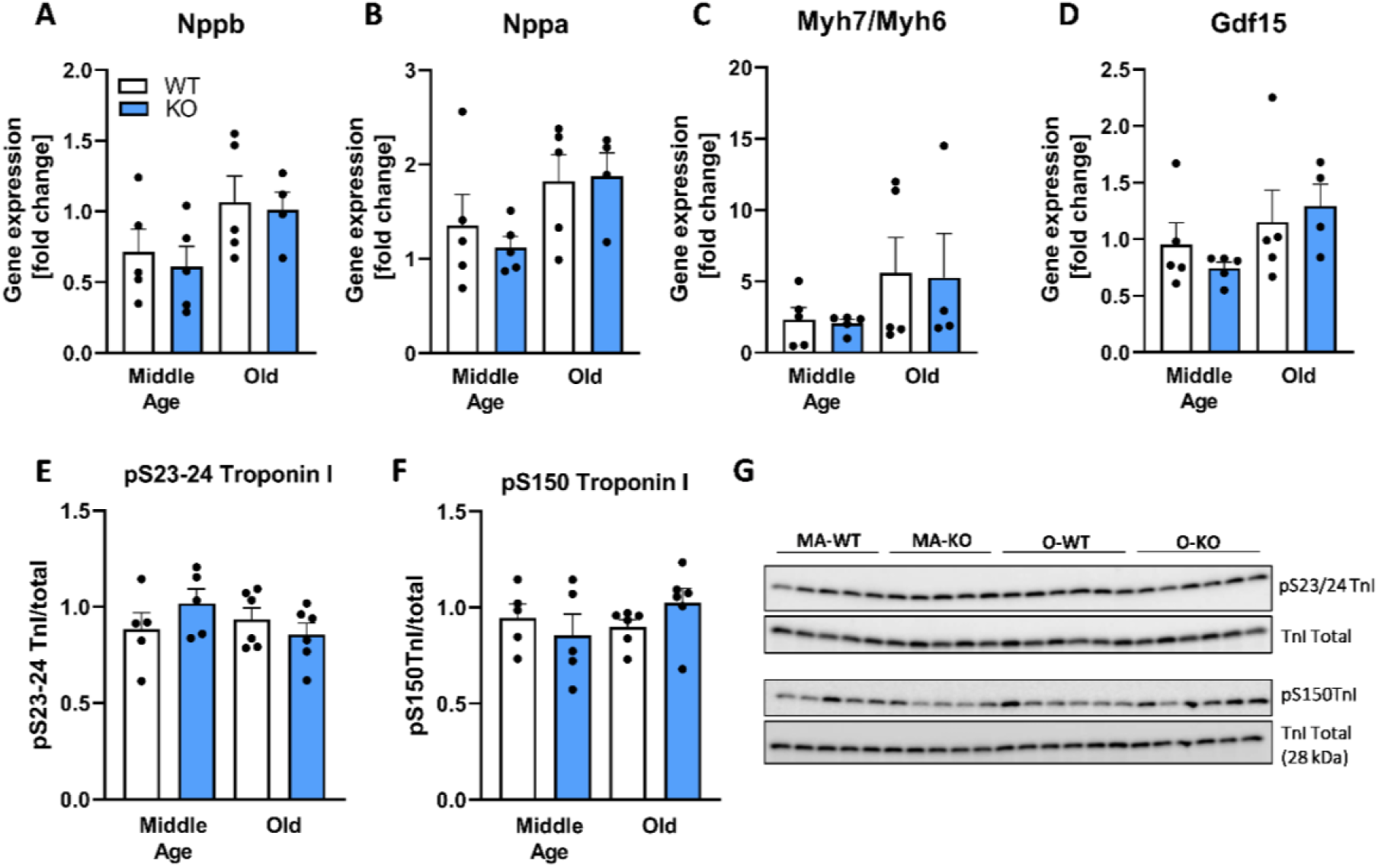
4EBP1 deletion showed no significant impact on gene expression levels of heart failure biomarkers and phosphorylation of myofilament protein Troponin I, regardless of age. **(A-D)** Transcript levels of heart failure markers (Nppb, Nppa, Myh7/Myh6 and Gdf15) are not different between 4EBP1 KO and WT mice regardless of age. However, the levels of **(B)** Nppa and **(D)** Gdf15 increase with age. Phosphorylation of **(E)** S23-24 and **(F)** S150 residues of Troponin I, which regulates cardiac contraction and relaxation, do not differ between WT and KO mice regardless of age. **(G)** Western blot of Troponin I phosphorylation in mouse heart. **(A-F)** Values represent means ± SEM. One-way ANOVA was performed, n=4-6.

Phosphorylation of myofilament proteins, e.g. Troponin I (TnI), regulates cardiac systolic and diastolic function [45]. Phosphorylation of TnI at S23/24 is reduced in failing hearts and results in slower myocardial relaxation, whereas phosphorylation at both S150 and S23/24 sites increase in myocardial ischemia [45, 46]. We measured TnI phosphorylation at S23/24 and S150 and phosphorylation of these residues in 4EBP1 KO hearts are similar to WT hearts (Fig. 3E-G). This suggests that phosphorylation of Troponin I is not involved in the decline of cardiac function in 4EBP1 KO mice.

### Deletion of 4EBP1 dysregulates protein homeostasis in mouse hearts

Since mTORC1/4EBP1 pathway is responsible for protein synthesis and is a key regulator of proteostasis, we examined whether dysregulated protein synthesis and degradation contribute to the accelerated cardiac aging observed in middle-aged 4EBP1 KO mouse hearts. We used stable isotope, deuterium oxide (D_2_O), labeling combined with GC-MS to measure bulk protein fractional synthesis rates (FSR) as an indication of protein turnover in various cellular fractions in middle-aged 4EBP1 KO and WT hearts. Bulk protein FSR of cytosolic, myofibrillar and mitochondrial fractions did not show any significant differences between middle-aged 4EBP1 KO and WT hearts (Supplemental Fig. 3A-C). However, middle-aged 4EBP1 KO mouse hearts have higher RNA synthesis compared to age-matched WT controls (Fig. 4A). Ribosomal RNA (rRNA) constitutes majority of pooled RNA and ribosomal biogenesis is a result of coordinated synthesis of rRNAs and proteins [47]. These results suggest higher ribosomal biogenesis in 4EBP1 KO mouse hearts. Moreover, we observed a trend of increased synthesis of collagen in middle-aged 4EBP1 KO hearts compared to WT controls (p=0.0791, Fig. 4B).

**Figure 4.**
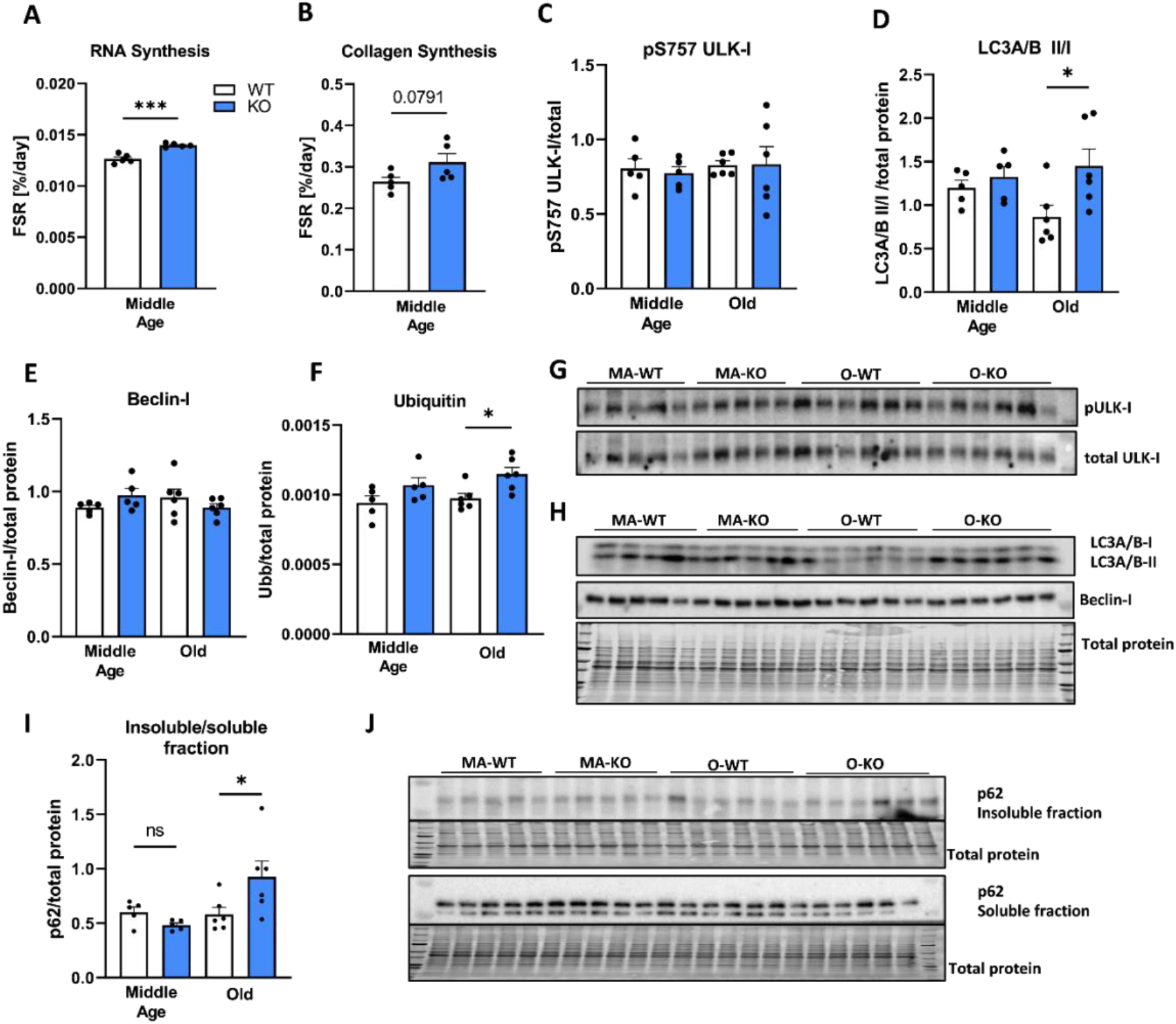
Proteostasis is dysregulated in 4EBP1 KO mouse hearts. Middle-aged 4EBP1 KO mouse hearts showed **(A)** greater RNA and **(B)** collagen synthesis. **(C)** ULK-I phosphorylation is not affected by 4EBP1 deletion. **(D)** Old 4EBP1 deficient mice have significantly increased protein ubiquitination compared to WT mice. **(E)** LC3A/B II/I ratio is significantly higher in old 4EBP1 KO hearts. **(F)** Protein expression of Beclin-I is not changed in 4EBP1 KO mouse hearts comparing to WT regardless of age. **(G)** Western blot of ULK-I phosphorylation in mouse heart. **(H)** Western blot of LC3A/B I and II and Beclin-I expression in mouse heart. **(I)** p62 expression is higher in insoluble fraction of old 4EBP1 KO mouse heart compared to WT **(J)** Western blot of p62 expression levels in soluble and insoluble fraction. Values represent means ± SEM. **(A, B)** Unpaired two-tailed Students t-test and **(C-F)** one-way ANOVA were performed, n=5-6. *p< 0.05; ***p< 0.0001

ULK-I is a mTORC1 downstream effector responsible for regulating protein degradation by autophagy [32, 48]. Phosphorylation of S757 residue (a target site of mTORC1) of ULK-I does not change due to 4EBP1 deficiency (Fig. 4C, G). To further assess protein degradation processes we examined changes in autophagy-associated proteins, Beclin-I, LC3 and p62. Autophagy marker LC3-II/I ratio increases in old 4EBP1 KO mouse hearts compared to old WT controls (Fig. 4D, H) but is not different between middle-aged WT and 4EBP1 KO mice (Fig. 4D, H). Simultaneously expression levels of Beclin-I, a protein involved in autophagy initiation process, are not different between WT and 4EBP1 KO mice (Fig. 4E, H). Old 4EBP1 KO mouse hearts show significantly higher protein ubiquitination compared to WT mice (Fig. 4F, Supplemental Fig. 3F). Middle-aged 4EBP1 KO mice show a trend of higher protein ubiquitination in comparison to WT (Fig. 4F). Interestingly, we observed that levels of p62, a cargo receptor for ubiquitinated proteins to target them for degradation, increases in the insoluble protein fractions of old 4EBP1 KO compared to old WT hearts (Fig. 4I, J). p62 accumulates when autophagy is inhibited, which can suggest dysfunctional autophagosome degradation. Moreover, 4-hydroxynonenal (4-HNE), marker of lipid peroxidation and oxidative stress which can induce autophagy [49] is higher in old hearts when compared to middle-aged hearts, but it is not different between WT and 4EBP1 KO mice (Supplemental Fig. 3D-E). Taken together, these data suggest that hyperactivation of mTORC1/4EBP1 downstream signaling pathway dysregulates protein degradation and proteostasis in the heart at advanced ages.

## Discussion

mTORC1 is essential for embryonic cardiovascular development and cardiac [50]. However, chronically elevated mTORC1 signaling also contributes to cardiac diseases and aging while inhibition of mTORC1 by rapamycin treatment reverses age-related cardiac dysfunction and hypertrophy in old mice [20, 34, 51]. Previous study on *Drosophila* has shown that muscle-specific overexpression of 4EBP1 increased lifespan and contributed to improved proteostasis [35, 52]. With regard to cardiac function, d4EBP null mutant flies show stress-induce failure rate of cardiac function [36]. Conversely, cardiac specific overexpression of d4EBP significantly reduced age-related decline in cardiac function [36]. However, the role of mTORC1/4EBP1 axis in cardiac aging has not been established in mammalian models. Using a 4EBP1 KO mouse model, we demonstrated that hyperactivation of mTORC1/4EBP1/eIF4E axis exacerbates cardiac dysfunction in middle-aged and old hearts independently of cardiac hypertrophy and without affecting function in young hearts. We observed that young WT and 4EBP1 KO male mice have similar cardiac function based on all measured cardiac function parameters (Fig. 1). However, middle-aged 4EBP1 KO mice show worse diastolic and systolic function than middle-aged WT mice (Fig. 2A and C). This dysfunction further declines in old age and old 4EBP1 KO mice also have worse cardiac function than old WT mice (Figure 2A and C). These results suggest that 4EBP1 deletion, which mimics hyperactive mTORC1/4EBP1 axis, accelerates cardiac aging in mice. Therefore, these findings indicate that mTORC1/4EBP1 signaling pathway plays a significant role in mediating age-related decline of cardiac function and are consistent with previous studies on *Drosophila*.

We detected no difference in gene expression of the cardiac hypertrophy (Nppb, Nppa, Myh7/Myh6, Gdf15), cellular stress (Atf4) or cellular senescence (Il6, p16, p19) biomarkers between 4EBP1 KO and WT mice, regardless of age (Fig. 3A-D). These results suggest that these mechanisms likely do not contribute to the worsen cardiac function caused by deletion of 4EBP1. We also investigated phosphorylation of myofibrillar protein, Troponin I, which plays an important role in cardiac contraction and relaxation [53]. We found that phosphorylation of Troponin I at residues S150 and S23/24 do not change due to deletion of 4EBP1, which indicates that hyperactive mTORC1/4EBP1 axis does not impair cardiac function through alteration of TnI phosphorylation.

Proteostasis imbalance is a key factor contributing to decline of cardiac function and aging. Accumulation of either abnormal or misfolded proteins as aggregates indicates a lack of proteostatic maintenance. Hyperactivation of mTOR signaling may result in impaired proteostatic maintenance and excessive amount of protein aggregates and misfolding that ultimately lead to cellular dysfunction [54]. In contrast, inhibition of mTORC1 either by CR or rapamycin have been shown to improve proteostasis [19]. Moreover, Demontis et al. have shown that FOXO/4EBP signaling regulates proteostasis in *Drosophila* muscles during aging [52]. Because mTORC1 signaling is a major regulator of proteostasis we utilized deuterium oxide labeling method to elucidate the effect of hyperactivation of mTORC1/4EBP1 axis on ribosomal biogenesis and protein synthesis. We showed that ribosomal biogenesis, is significantly increased in middle-aged 4EBP1 KO mice comparing to their WT littermates. mTORC1 regulates ribosome biogenesis by various mechanisms and it is well established that ribosomal proteins play a key role in protein synthesis thus contributing to proteostasis maintenance [55]. Steffen et al. have shown that reduced ribosomal biogenesis mediated by inhibition of TOR signaling is responsible for beneficial effects of CR in *S. cerevisiae* [56]. The observed increase in ribosomal biogenesis in middle-aged 4EBP1 KO hearts suggests that mTORC1/4EBP1 hyperactivation promotes ribosomal biogenesis.

In terms of protein synthesis, we observe a near-significant increase in collagen synthesis. Surprisingly, in light of increased ribosomal biogenesis we observed no difference in protein synthesis in cytosolic, myofibrillar or mitochondrial fractions. Although, we did not observe any differences in bulk protein synthesis in different subcellular fractions, we detected significant increase in protein ubiquitination in old 4EBP1 KO and a trend towards higher levels of ubiquitination in middle-aged 4EBP1 KO mouse hearts compared to age-matched WT hearts. This suggests that hyperactivated mTORC1/4EBP1 axis, may result in misfolded or aggregation-prone proteins and ultimately leading to accumulation of non-functional proteins which are marked for degradation. We do not observe differences in phosphorylation of ULK-I at the mTORC1 target site (S757) or the expression of Beclin-I between 4EBP1 KO and WT mice. This suggests that neither mTORC1/ULK-I signaling pathway nor autophagy initiation process are altered due to 4EBP1 deletion. However, we observed higher LC3-II/I ratio and accumulation of p62 in old 4EBP1 KO mice compared to WT which suggests accumulation of autophagosome and impaired autophagy mechanism. In conclusion, these findings suggest that proteostasis in 4EBP1 KO mice is dysregulated due to impaired autophagy and it potentially contributes to the accelerated age-related decline in cardiac function observed in 4EBP1 KO mice.

A limitation of this study is that we used a whole body 4EBP1 KO mouse model. Deficiency of 4EBP1 in other organs may have indirect impacts on the heart. We primarily used male mice for the study and observed no sex-specific differences in the parameters that we assessed in both male and female mice. However, we cannot exclude sex-specific changes in the parameters we did not study in female mice. Another limitation is that single time point (15 days) labeling protocol may not reveal changes in the dynamic pool size. The differences in synthesis rates may therefore reflect the fraction of the pool that is turning over [57]. A timecourse experiment would differentiate the effect of changes in pool size versus synthesis rates.

Taken together, our study supports an important role of mTORC1/4EBP1 signaling in cardiac aging and suggests that dysregulated proteostasis induced by hyperactive mTORC1/4EBP1 axis, in part, contributes to the deterioration of cardiac dysfunction with age. Further investigation of mTORC1/4EBP1/eIF4E signaling and other branches of mTORC1 signaling axis in the aging heart is needed to further dissect the molecular mechanisms involved in the beneficial effects of mTORC1 inhibition in cardiac aging. A better understanding of mTORC1/4EBP1 signaling pathway and mechanisms involved in cardiac aging will facilitate the development of novel and effective therapies with minimized off-target effects.

## Acknowledgements

We thank Dr. Nahum Sonenberg at McGill University for generously providing the 4EBP1 KO mouse line for this study. We acknowledge the funding support from NIH R00AG051735 (YAC) and from AHA 23POST1012408 (KAK).

## Author Contribution

W.Z., B.F.M. and Y.A.C. designed the study. W.Z., K.A.K., C.J.K., F.F.P.III, B.F.M. and Y.A.C. acquired, analyzed, and interpreted the data. W.Z. and Y.A.C. wrote the manuscript. All authors contributed to the review and revision of the manuscript.

## Disclosures

W.Z., K.A.K., C.J.K., F.F.P.III, B.F.M. and Y.A.C. declare no competing interests.

## Supplemental Figures

**Supplement Figure 1.**
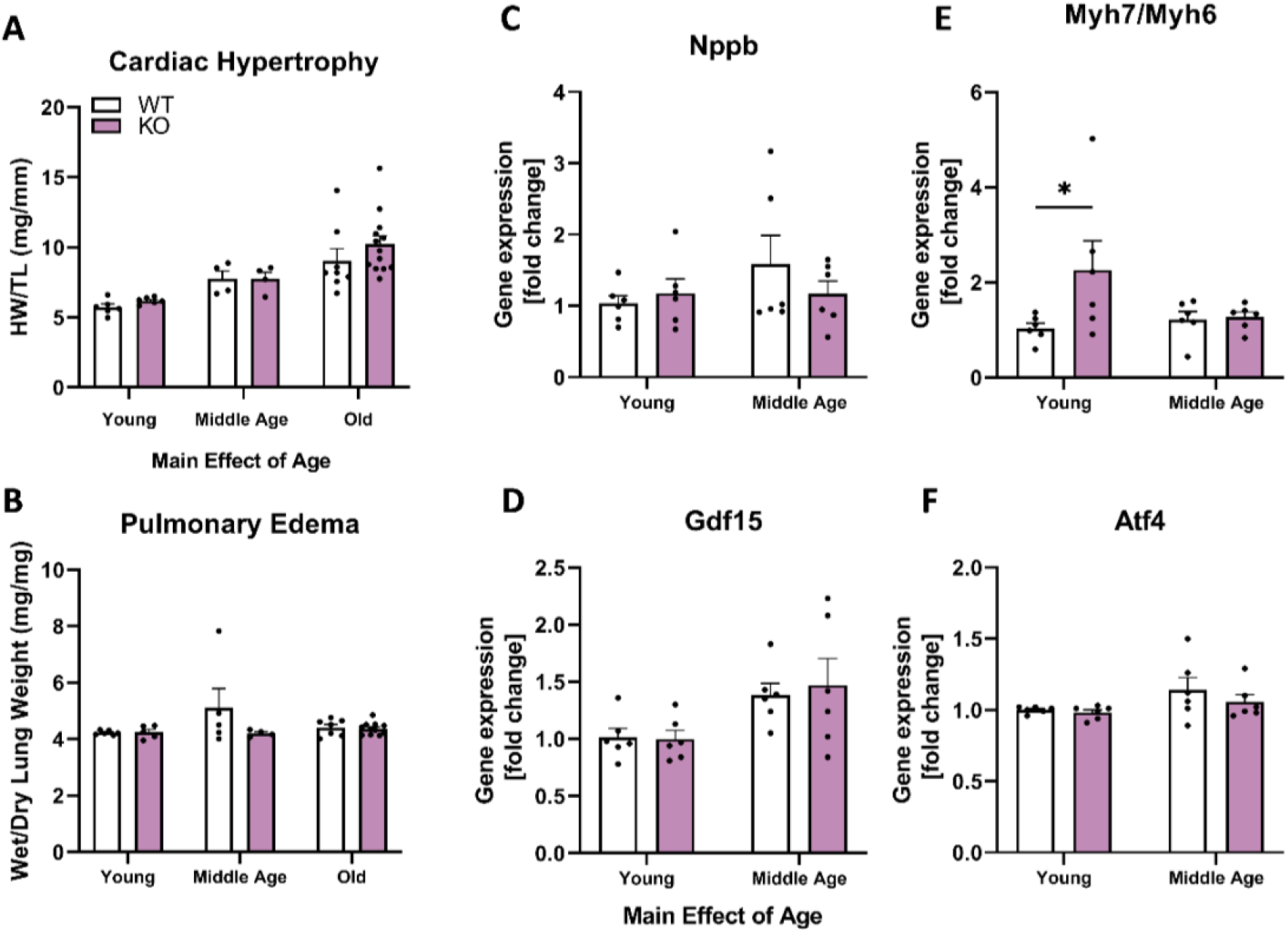
Deletion of 4EBP1 protein does not have an impact cardiac hypertrophy and pulmonary edema or gene expression levels of cardiac hypertrophy and cellular stress biomarkers in 4EBP1 KO female mice compared to WT. **(A, B)** Cardiac hypertrophy in female mice increases with age, however KO mice showed no altered hypertrophy comparing to WT mice, similarly 4EBP1 KO female mice do not have pulmonary edema **(C-F)** Gene expression levels of heart failure markers (Nppb, Myh7/Myh6, Gdf15 and Atf4) are not different between 4EBP1 KO and WT mice regardless of age. **(D)** However, expression levels of Gdf15 shows a significant increase with age regardless of genotype. Values represent means ± SEM. Two-way ANOVA was performed, n=4-6

**Supplement Figure 2.**
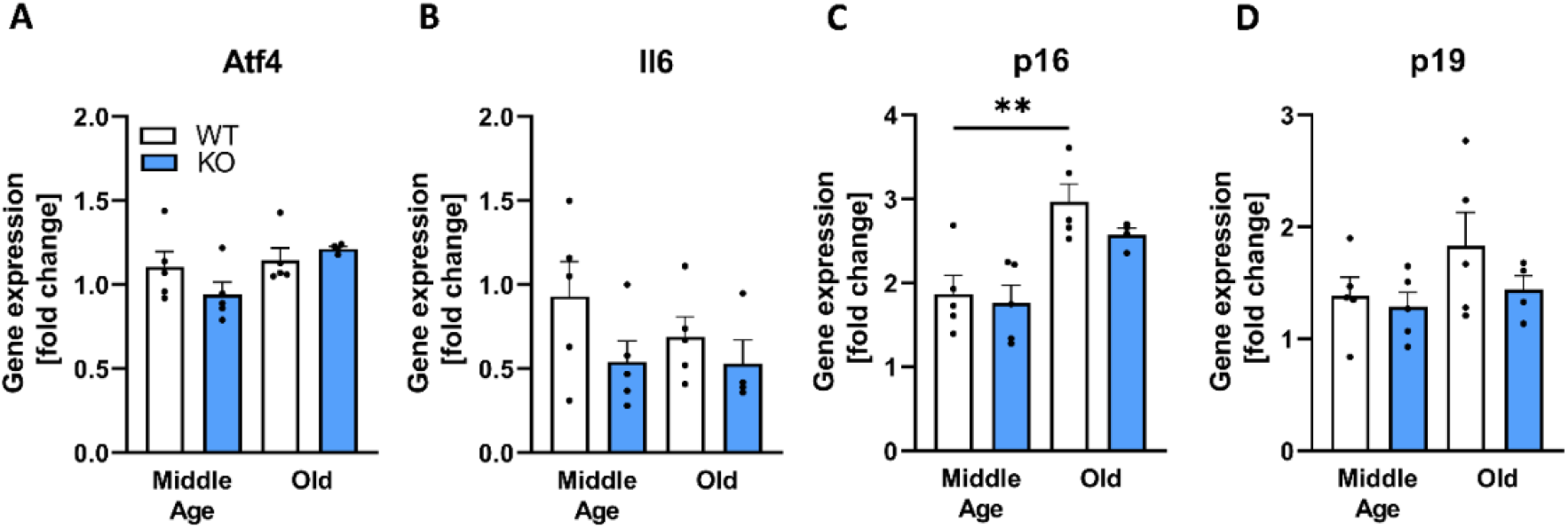
Complete 4EBP1 deficiency does alter gene expression levels of cellular stress and senescence biomarker in middle-aged and old male mice. **(A)** Gene expression levels of cellular ER stress marker (Atf4) are not different between 4EBP1 KO and WT mice regardless of age. **(B-D)** Gene expression levels of cellular senesces markers (IL6, p16, p19) are not different between 4EBP1 KO and WT male mice. Values represent means ± SEM. One-way ANOVA was performed, n=5.

**Supplement Figure 3.**
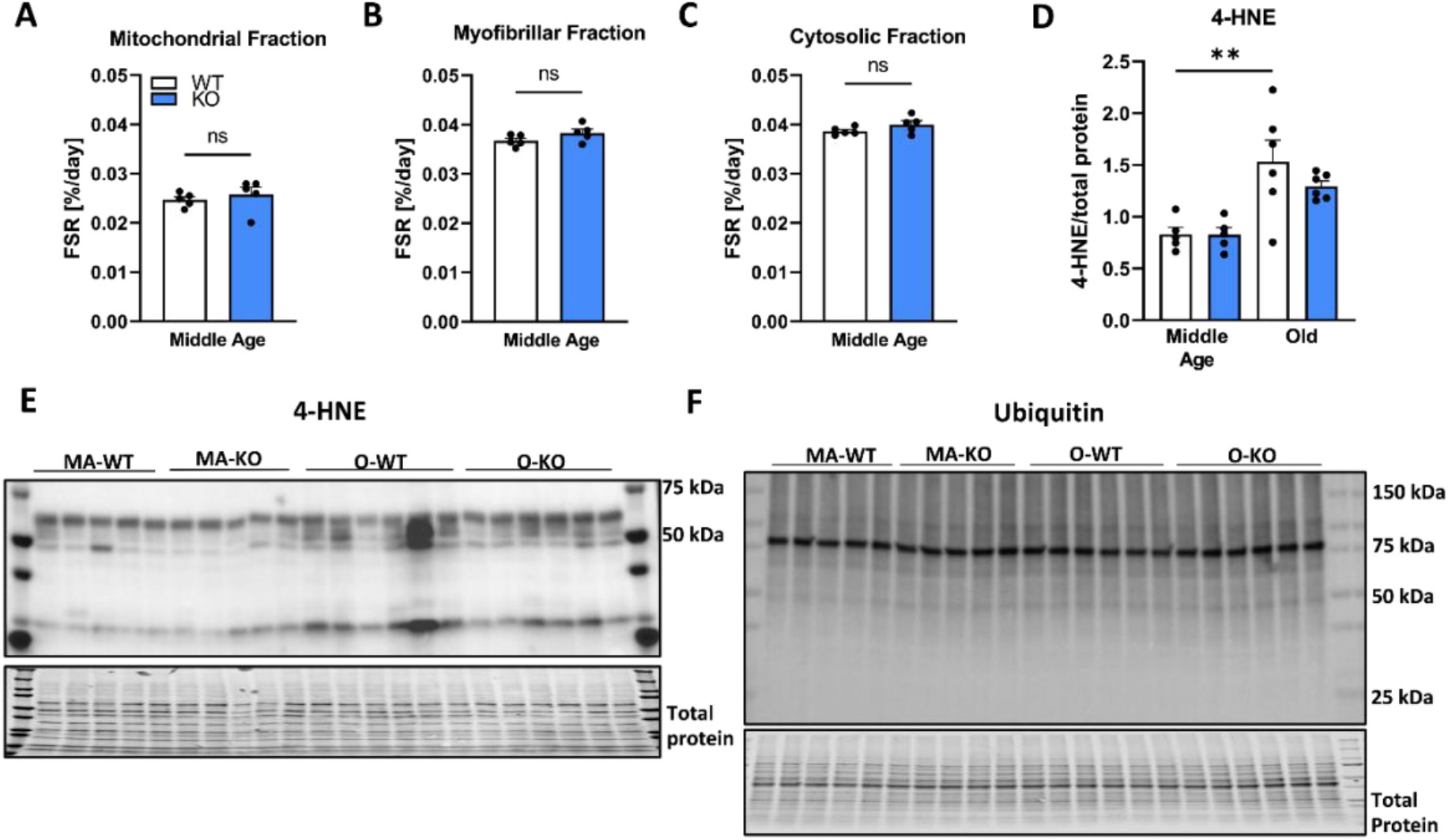
Protein synthesis and marker of oxidative stress are unchanged due to deletion of 4EBP1 but it increases protein ubiquitination. **(A-C)** Synthesis of mitochondrial, myofibrillar and cytosolic proteins is not different in 4EBP1 KO mice comparing to WT. **(D)** 4-HNE content in middle age and old WT and 4EBP1 KO mouse hearts. **(E)** WB image of 4-HNE. **(F)** WB image of protein ubiquitination in middle-aged and old WT and 4EBP1 KO mouse heart. Values represent means ± SEM. **(A-C)** Unpaired two-tailed Student’s t-test was performed, n=5. **(D)** One-way ANOVA was performed, n=5-6.

## Notes

### Competing Interest Statement

The authors have declared no competing interest.

